# HeatChips: A versatile, low-cost and microscopy-compatible heating system for microfluidic devices

**DOI:** 10.1101/2022.11.15.516605

**Authors:** Théo Aspert, Gilles Charvin

## Abstract

Microfluidic systems are widely used in biology for their ability to control environmental parameters. Specifically, cell culture or chemistry in microfluidic devices requires tight control of the temperature. In addition, microfluidic devices can be made transparent to visible light and compatible with inverted microscopes. Yet, the current temperature control systems that allow high-resolution microscopy either require a set of complex secondary channels, a bulky, expensive, and microscope-dependent incubator, or fail to produce a homogenous temperature profile across the sample area. Here, we present HeatChips, a simple, cost-effective system to heat samples inside PDMS-based microfluidic devices in a homogeneous manner. It is based on a transparent heating glass in contact with the top of the microfluidic device, and a contactless, infrared temperature sensor attached to the objective that directly reads the temperature of the bottom of the chip. This portable system is compatible with most chip designs and allows imaging of the sample on inverted microscopes for extended periods of time without any optical restriction, for a cost of less than 100€.

**Specifications table:** 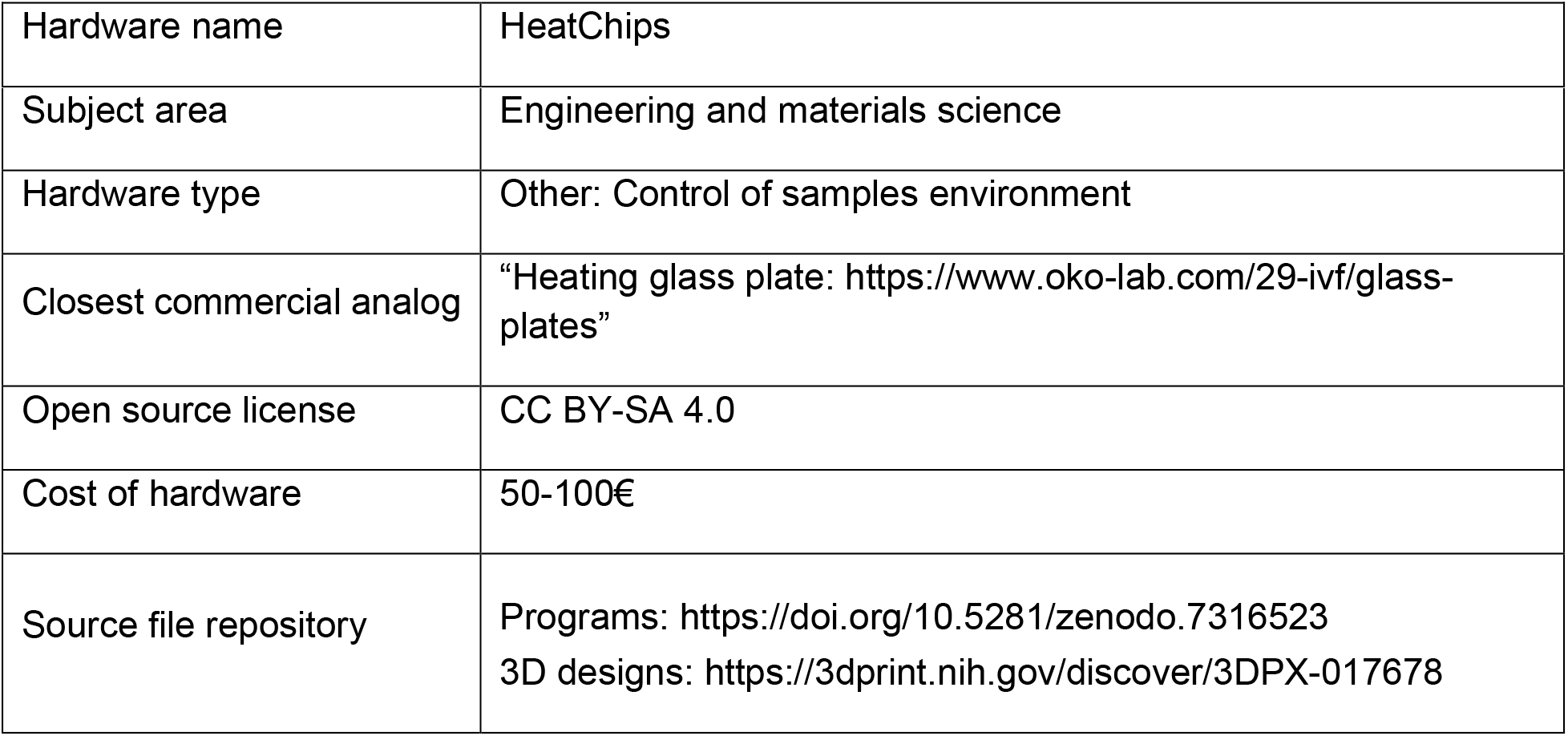

## 1. Hardware in context

Temperature is a critical parameter for most laboratory experiments, may it be for physical, chemical, or biological applications. In particular, since a major asset of microfluidic technology is to offer tight control of the environment of a sample of interest, the temperature also often needs to be controlled in these experiments [1].

To this end, a variety of strategies have been developed, with different levels of integration, range of temperature, accuracy, homogeneity, and response time. Some of them use external heaters such as light or acoustic waves directed toward the device [2–4], while other approaches consist of flowing a pre-heated liquid in secondary channels of the device [5] (exhaustive review in [1]).

However, the available solutions are restricted when the microfluidic device needs to be imaged for extended periods of time and for general applications in order to not block the optical path or interfere with image acquisition. In this case, two types of solutions prevail: to heat the environment of the device to the desired temperature or to directly heat the device by contact with a hot source.

For example, stagetop incubators or incubators surrounding the microscope offer a homogeneous temperature across the whole device and control of gas composition. However, this involves a voluminous and expensive piece of equipment (>15k€ for commercial solutions [6]) that often comes as an add-on unit for a given microscope. While Do-it-yourself (DIY) solutions have been recently proposed for a cost of around 350€ [7], this equipment remains cumbersome, difficult to reproduce, and microscope-dependent.

The other existing solution is to use the sample holder as a hot source [8–10], which would in turn heat the device by diffusion. This inexpensive solution might however lead to a gradient of temperature between the edges of the device, in contact with the heat source, and the center, in contact with the air because of the aperture of the sample holder. This inhomogeneity can be reduced by adding a complementary heating source from the objective if using oil objectives with samples with a limited surface area to scan [8], but this remains specific cases of applications. Moreover, to enslave the sample temperature to the setpoint, these two types of solutions measure the temperature of the heating source (namely, the air or the sample holder) instead of a direct measurement of the sample’s temperature, which can lead to a static error between the setpoint and the actual sample’s temperature.

Alternatively, mounting the microfluidic device onto a heating glass allows homogeneous, precise, and stable heating (for example, the heating glass from Okolab©, Italy). Indeed, these glasses are coated on one side with a thin layer (*i.e.* 200nm thick) of conductive material (typically, Indium Tin Oxide or (ITO)) that, when connected to a power supply, heats up by the Joule effect. Since a temperature sensor can be inserted into the glass, and since the sample of interest is in direct contact with the heat source, the measured temperature is therefore very close to that of the sample. In addition, this solution is compatible with imaging since the glass remains transparent. Yet, such a solution requires a high working distance since it involves placing a 0.5mm thick heating glass in between the sample and the objective. Consequently, it is incompatible with most high-magnification objectives and negatively affects the resolution of the image. Moreover, these commercially available devices are specific to each microscope, which limits their versatility.

Here, we describe a heating system that is compatible with most microfluidic devices, microscopes, and objectives, without affecting the image quality, while allowing to precisely maintain a homogeneous temperature along the sample area for extended periods of time, in a cost-effective and easily reproducible manner. The system is based on a heating, transparent glass that can be put in contact with the top of PDMS-based microfluidic devices. An infrared probe facing the glass slide of the device is used to directly read the temperature of the sample, and the whole system is controlled with a simple, Arduino-based electronic circuit and program (Figure 1).

**Fig. 1.**
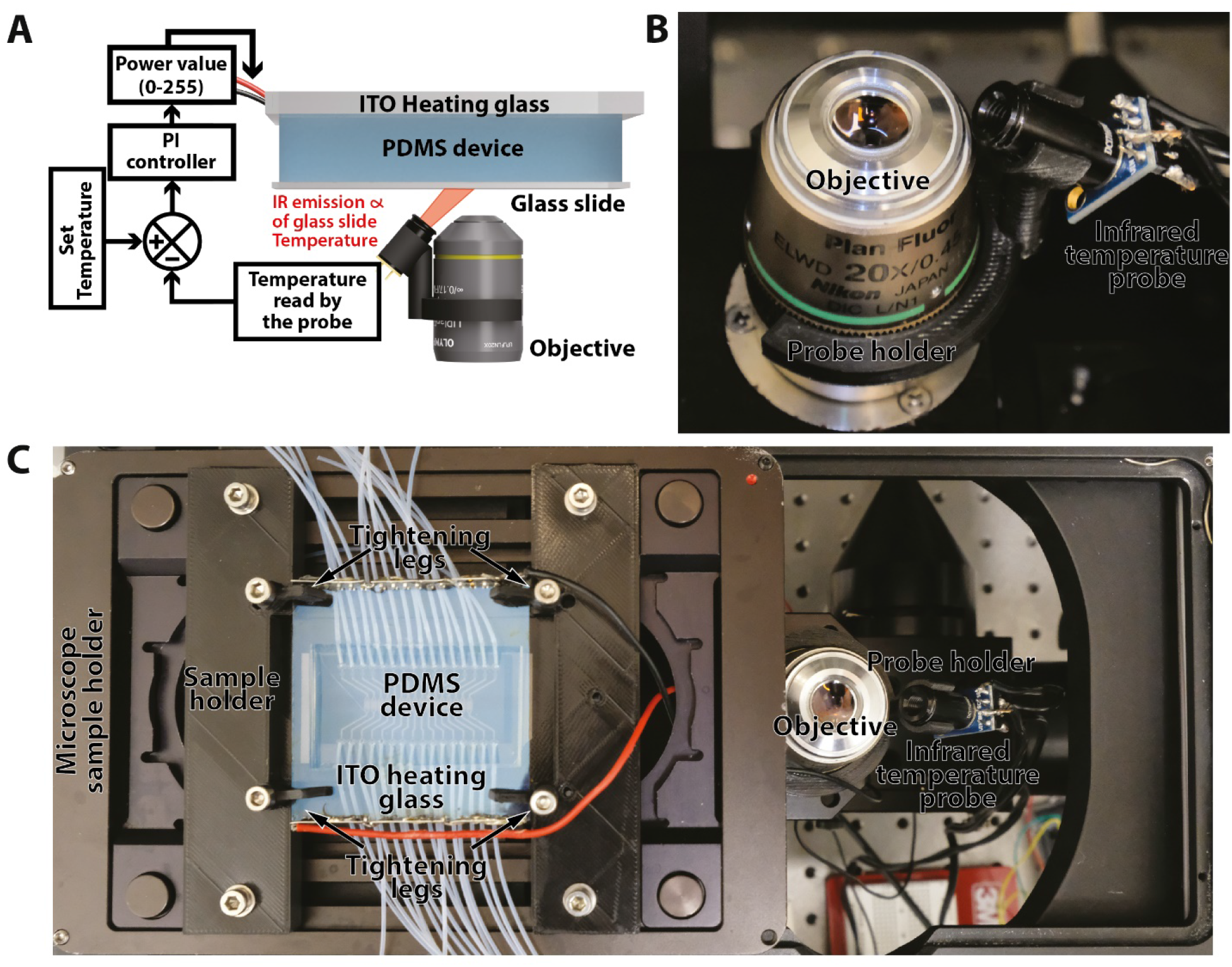
(A) Scheme of the operating principle of HeatChips. The scale of the objects is not representative. (B) Picture of the infrared temperature probe mounted on an objective. (C) Picture of the top view of the HeatChips setup, with the microscope sample holder set aside the microscope for better visualization.

## 2. Hardware description

### Hardware composition

As any heating system, HeatChips is composed of two parts: a heating part, and a control part.

#### Heating part

HeatChips was designed to heat a sample inside a PDMS device, in a homogeneous and compact way, while being compatible with microscopy. Therefore, we decided to use an ITO-coated glass (see the previous section for details) as a heat source, in direct contact with the top of the microfluidic device (Figure 1A). Therefore, the heating system is independent of the microscope configuration, with equal temperature distribution on the plane of the sample (located on the glass slide), and is fully compatible with inverted microscopes. Indeed, as mentioned in 1., an ITO-coated glass is transparent to visible light, hence it does not affect the brightfield illumination, nor epifluorescence excitation since it is placed after the sample on the optical path (Figure 1A).

#### Control part

The control part is made of a probe measuring the temperature of the sample, connected to an electronic system controlling the power sent to the *ITO glass.*

##### Temperature probe

To make the temperature measurement independent of any external factors that may vary from one experimental setup to another (room temperature, materials in contact with the device, thickness of the device, …), we used a temperature probe that directly measures the temperature of the glass slide, which is only a few tens of microns distant from the sample.

The temperature probe is an infrared sensor attached to the objective using a 3D-printed holder, pointing toward the sample and measuring the temperature of the glass slide without contact with the sample. Indeed, the probe contains a 5.5μm-14μm pass-band filter, wavelengths to which the crosslinked PDMS is opaque [11,12], allowing it to block most of the photons emitted by the heating glass in this range. We tested that experimentally by measuring the output of the probe pointing toward either: the wall, the glass slide, the glass slide bound to a 5mm thick PDMS block (1:10 curing agent ratio), a 5mm thick PDMS block, the *ITO glass* receiving 1W of energy, or a combination of these objects (Figure 2A). If the resulting temperature from the ITO was around ~37°C, the temperature read with the probe pointing toward the glass slide + ITO glass was attenuated down to ~25°C, suggesting that the glass slide is not transmitting most of the emissions from the 5.5μm – 14μm spectrum (Figure 2B). Similarly, the temperature measured in the configuration PDMS + ITO glass was indistinguishable from that of the PDMS alone, demonstrating that the PDMS acts as a screen for this spectrum of emission. Altogether, this suggests that the *infrared temperature probe* only measures the temperature of the surface of the PDMS and of the *glass slide,* when pointing toward the *glass slide* side of a PDMS-based microfluidic device.

**Fig. 2.**
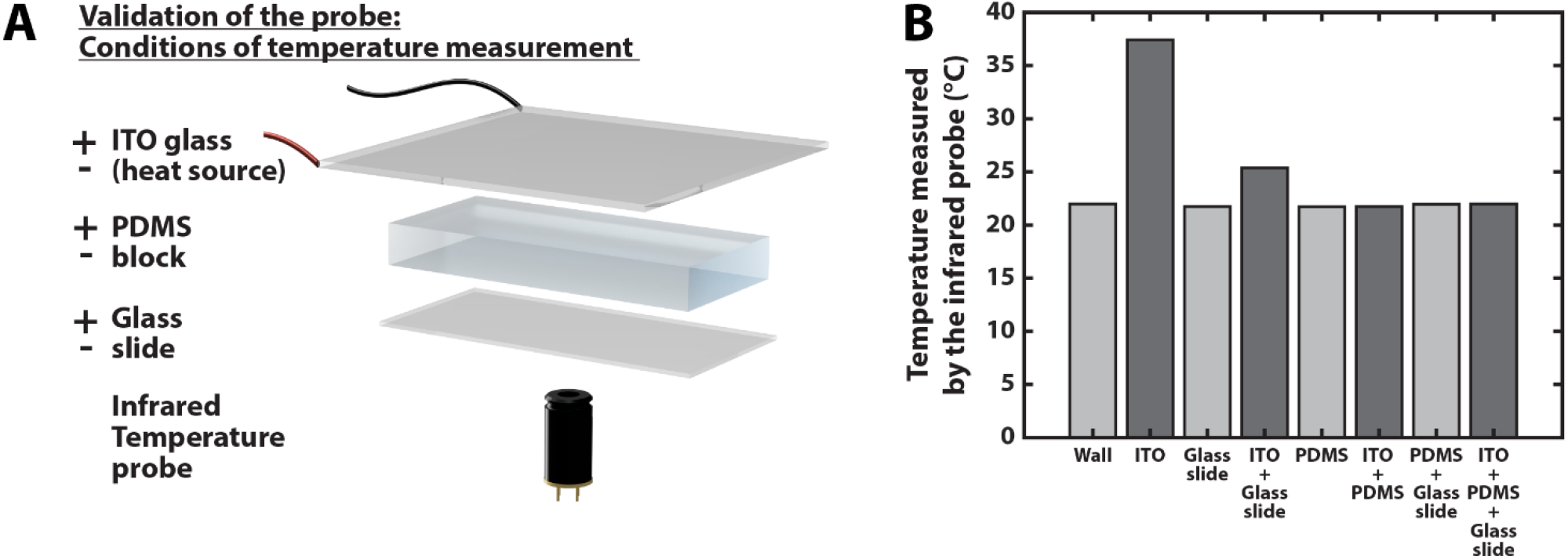
(A) Representation of the different conditions used to determine the validity of the infrared probe, in front of which a glass slide, a 5mm-thick PDMS block, or an ITO glass receiving 1W of electrical power, were present (+) or not (-). (B) Bar plot representing the temperature measured by the infrared probe in the conditions defined in (A).

##### Arduino-based electric circuit

To control the temperature of the sample, we developed a simple feedback loop system between the temperature from the probe and the output power sent to the heating glass. It is based on a proportional-integral-derivative (P-I-D) controller computed by an Arduino card reading the temperature probe, and sending a Power Width Modulation (PWM) signal to a MOSFET connected to the output power, a 5V 1A phone charger (Figures 1 and 5).

### Range of application

This heating system has been specifically designed to maintain samples (cells, fluids, …) inside PDMS-based devices at a constant temperature above room temperature in order to image them under an inverted microscope for extended periods of time.

Furthermore, heating from the sample holder usually requires an aperture of restricted size in order to reduce the gradient of temperature between the center of the device and the edges, in contact with the heat source. Here, since the heat source comes from the top of the device, the aperture of the sample holder can almost be as large as the device itself. Therefore, HeatChips is particularly useful to scan large areas of a microfluidic device.

Moreover, it is particularly suitable for experiments where precise control of the air composition is not required, as this device can then replace a bulky and expensive incubation chamber placed around the microscope.

In theory, HeatChips works with any PDMS (or any elastomer that is opaque to the 5.5μm - 14μm spectrum) based devices and is adaptable to most inverted imaging systems with minimal work (adjustments of the 3D printed designs).

The range of working temperatures is determined by the infrared temperature probe. The heating temperature is between ambient temperature and 60°C, and the system has maximum accuracy when working with an ambient temperature between 0 and 60°C.

#### Limitations

HeatChips is not suited for fast temperature switches or oscillatory patterns since the equilibrium time is reached between 10 and 30min (depending on the PDMS thickness, setpoint temperature, and ambient temperature, more information about the system’s characteristics in section 7. and in Figure 7).

Since HeatChips heats the sample from the top and by contact, there must be no obstruction between the glass and the top of the PDMS. Thus, leveling the surface on which the PDMS is cured can improve the flatness of the surface and ensure homogeneous heating of the sample. Moreover, the glass might need to be cut or drilled (using standard glasswork methods), or tubing holes can be punched from the sides of the chip instead of from the top, in order for the tubing to not alter the contact.

## 3. Design files summary

| Design file name | File type | Open source license | Location of the file |
| --- | --- | --- | --- |
| Electronic diagram | Image | CC BY-SA 4.0 | Available with the article (Figure 5) |
| Sample holder | 3D files (.stl and .f3d) | CC BY-SA 4.0 | https://3dprint.nih.gov/discover/3DPX-017678 |
| Tightening legs | 3D files (.stl and .f3d) | CC BY-SA 4.0 | https://3dprint.nih.gov/discover/3DPX-017678 |
| Probe holder | 3D files (.stl and .f3d) | CC BY-SA 4.0 | https://3dprint.nih.gov/discover/3DPX-017678 |
| Arduino program | .ino text file | CC BY-SA 4.0 | https://doi.org/10.5281/zenodo.7316523 and https://github.com/TAspert/HeatChips |
| (Optional) read_HeatChips.m | .m text file (MATLAB) | CC BY-SA 4.0 | https://doi.org/10.5281/zenodo.7316523 and https://github.com/TAspert/HeatChips |
| Arduino Software | Free software from Arduino.cc | GNU GPL | https://www.arduino.cc/en/software |

<u>Electrical diagram</u>: the electronic design used for the control part of the heating system is depicted in figure 7.
<u>Sample holder</u>: 3D files relative to the 3D printed sample holder (depicted on figure 3). This part is used to hold the microfluidic device and can be attached to a standard microscope slide holder.
<u>Tightening legs</u>: 3D files relative to the 3D printed tightening legs (depicted on figure 3). They are small plastic pieces used to hold the *ITO glass* against the PDMS, and hold the PDMS device in place onto the *sample holder.* Print in PLA or similar plastic.
<u>Probe holder</u>: 3D files relative to the 3D printed probe holder (depicted on figure 3). It is used to hold the *infrared temperature probe* onto the objective of the microscope. Different variants are already available, but the design can be adapted to fit specific needs.
<u>Arduino programs</u>: Program used to read the temperature from the *infrared probe* and compute an output power using a Proportional Integral Derivative controller. Two versions of the program exist: *HeatChips_Arduino_Program_withMatlabOutput.ino* which outputs temperature and power values that can be plotted using *read_HeatChips.m* (see next file) and a standalone version *HeatChips_Arduino_Program_standalone.ino.*
<u>read_HeatChips.m</u>: MATLAB script to read the temperature in real-time. To use with the *HeatChips_Arduino_Program_withMatlabOutput.ino* arduino program.
<u>Arduino software</u>: Official software from Arduino.cc used to upload the *Arduino program* into the *Arduino microcontroller,* and optionally, to visualize the temperatures and power output.

**Fig. 3.**
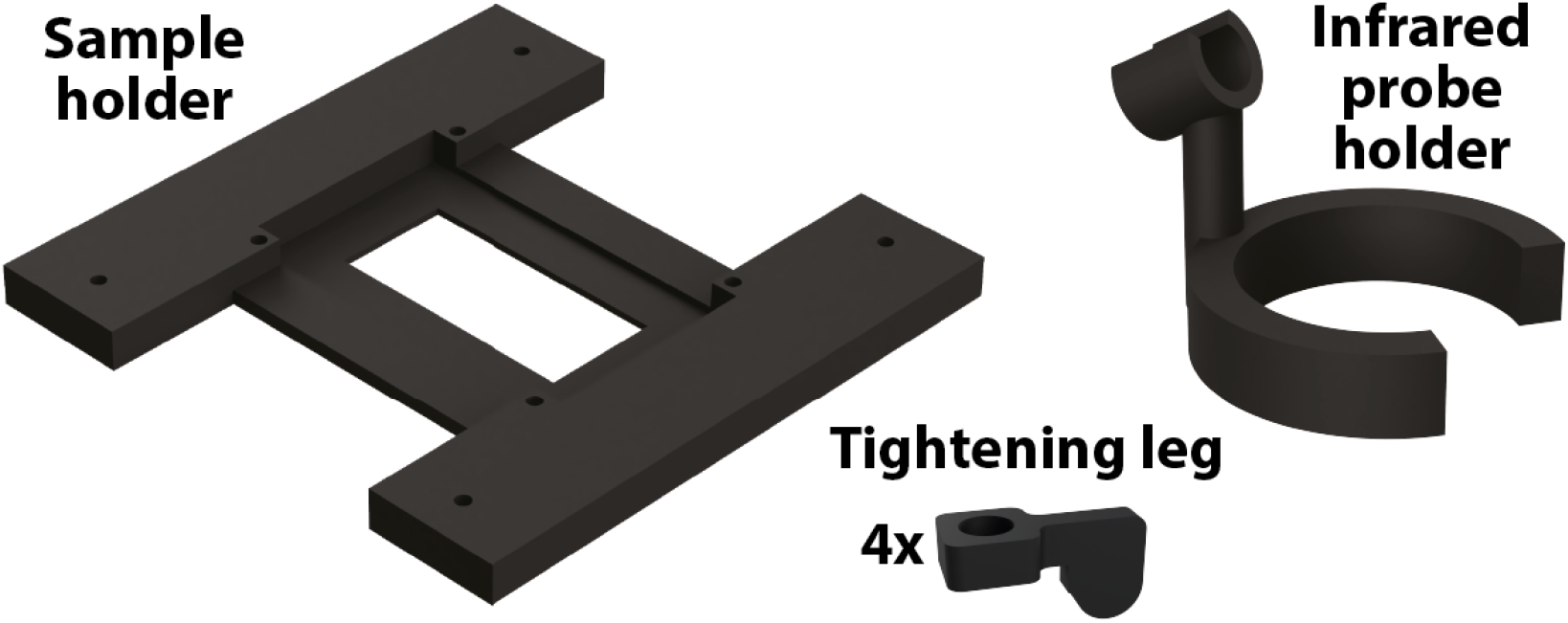
3D representation of the parts to be 3D printed.

## 4. Bill of materials summary

| Designator | Component | Qty | Cost per unit - € | Total cost - € | Source of materials |
| --- | --- | --- | --- | --- | --- |
| ITO glass | Indium Tin Oxide coated glass | 1 | 7.20 | 7.20 | https://t.ly/KpRp |
| Infrared temperature probe | GY-906, MLX90614ESF-DCI-000-SP | 1 | 30.03 | 30.03 | https://t.ly/8qey |
| Connection legs for the ITO glass | 10 legs for a 1mm thick glass | 2 | 0.68 | 1.36 | https://t.ly/y-Op |
| Arduino | Arduino microcontroller | 1 | 12.29 | 12.29 | https://t.ly/Ukfy |
| MOSFET | IRF520NPBF | 1 | 2.55 | 2.55 | https://t.ly/cx_O |
| 10kΩ Resistor | 10kΩ Resistor | 1 | 0.05-1.00 | 0.05-1.00 | https://t.ly/6EdR |
| Micro USB (or USB C) Port | Micro USB Port (or USB C) | 1 | 2.19 | 2.19 | https://t.ly/FS4_ |
| Micro USB (or USB C) power plug | Any 5V 1A or similar, micro USB or USB C power plug and cable | 1 | 5 € | 5 € | Widely available |
| Breadboard | Breadboard | 1 | 5.49 | 5.49 | https://t.ly/Zg5u0 |
| M3 screw and nut | M3 screw and nut | 8 | 0.1 | 0.8 | Any hardware store |

<u>ITO glass</u>: Indium Tin Oxide coated heating glass. For classical applications, use these specs: dimension 5ümm×5ümm, resistance: of the glass is ~13 Ω per cm^2^, ITO layer thickness: ~200nm.
<u>Infrared temperature probe</u>: Contactless temperature sensor based on the infrared emissions of the target.
<u>Connection legs</u>: (Optional) Small metallic parts to connect the ITO glass with electrical wires.
<u>Arduino</u>: Microcontroller driven by the *Arduino program* of Table 1. Component of the control electrical circuit.
<u>MOSFET</u>: Electronic component of the control part of the system.
<u>10kΩ resistor</u>: Electronic component of the control part of the system.
<u>Micro USB (or USB C) port</u>: Component bridging the control part and the heating part of the system.
<u>Micro USB (or USB C) power plug</u>: Power supply to plug on the port described above.
<u>Breadboard</u>: (Optional) Component to perform electrical tests of the reproduced circuit, or to replace a PCB.
<u>M3 screw and nut</u>: Used to assemble the *Tightening legs* to the *Sample holder* (and optionally, to lock the *Sample holder* with the microscope slide holder).

## 5. Build instructions

To reproduce HeatChips:

1. 3D print the *sample holder* (Figure 3 and Table 1). This part is used to hold and image the microfluidic device in place during a microscopy experiment, and to keep the heating glass in contact with the top of the device using four *tightening legs* of the desired size.
2. Clip the *sample holder* into the slide holder of the microscope to be used. (Optional) Lock the two parts together using 4× *M3 screws* and *bolts.*
3. 3D print four *tightening legs* (Figure 3 and Table 1).
4. 3D print the *probe holder* (Figure 3 and Table 1).
5. (Optional) Cut the *ITO glass* to the desired size, using a glass cutting tool, and drill holes at the desired position.
6. Clip the *connection legs* to two opposite sides of the *ITO glass*, and solder a cable to the legs (one cable to each of the two sides of the *ITO glass*) (Figure S1). The connection legs can be commercial (see Table 2), or homemade using small metallic clips or conductive tape. The goal of this component is to bring the electrical current to the ITO coating of the glass.
7. Solder 4 electrical cables to the *infrared probe* (Figure 4A, step 1).
8. Install the *infrared probe* into the *probe holder* (Figure 4A, step 2).
9. Clip the *infrared probe holder* onto the objective of the inverted microscope (Figure 4A, step 3).
10. Reproduce the electric circuit, using a breadboard or a PCB (Figure 7). Briefly:
  a. Solder 4 cables to the PCB pins of the infrared probe, and plug the cables into the associated Arduino I/O (GND on GND, VIN on 3V3 or 5V, SDA to Pin A4 and SCL to pin A5 (for an Arduino UNO, note that these pins might change from one Arduino model to another). The probe will then be able to send signals to the Arduino that can be interpreted by using the associated library (see code of the *Arduino program),* including the temperature of the target and the ambient temperature.
  b. Connect pin 11 (the pin number is defined in the *Arduino program*) to the gate of a *MOSFET* with a threshold voltage inferior to 5V, and to a *10kΩ* pull-up resistor.
  c. Connect the source of the *MOSFET* to the ground and the drain to one of the two cables of the *ITO glass*.
  d. Connect the other pole of the ITO glass to the VCC pin of the *micro USB* (or *USB C*) *port* and the GND pin of the port to the ground. Explanation of the circuit: The Arduino will compute an output number from the inputs from the temperature probe. This output number will be sent to pin 11 as a PWM signal between 0 and 5V, which will close or open the gate (resp.) of the MOSFET, hence modulating the average current passing through the heating glass, from the power supply.
11. Adjust the parameters of the *Arduino program* if needed (such as the Proportional and Integral coefficients, the setpoint, or the reading frequency, see comments in the code to see the effects of each parameter), and upload it to the Arduino card, using the *Arduino software.*

**Fig. 4.**
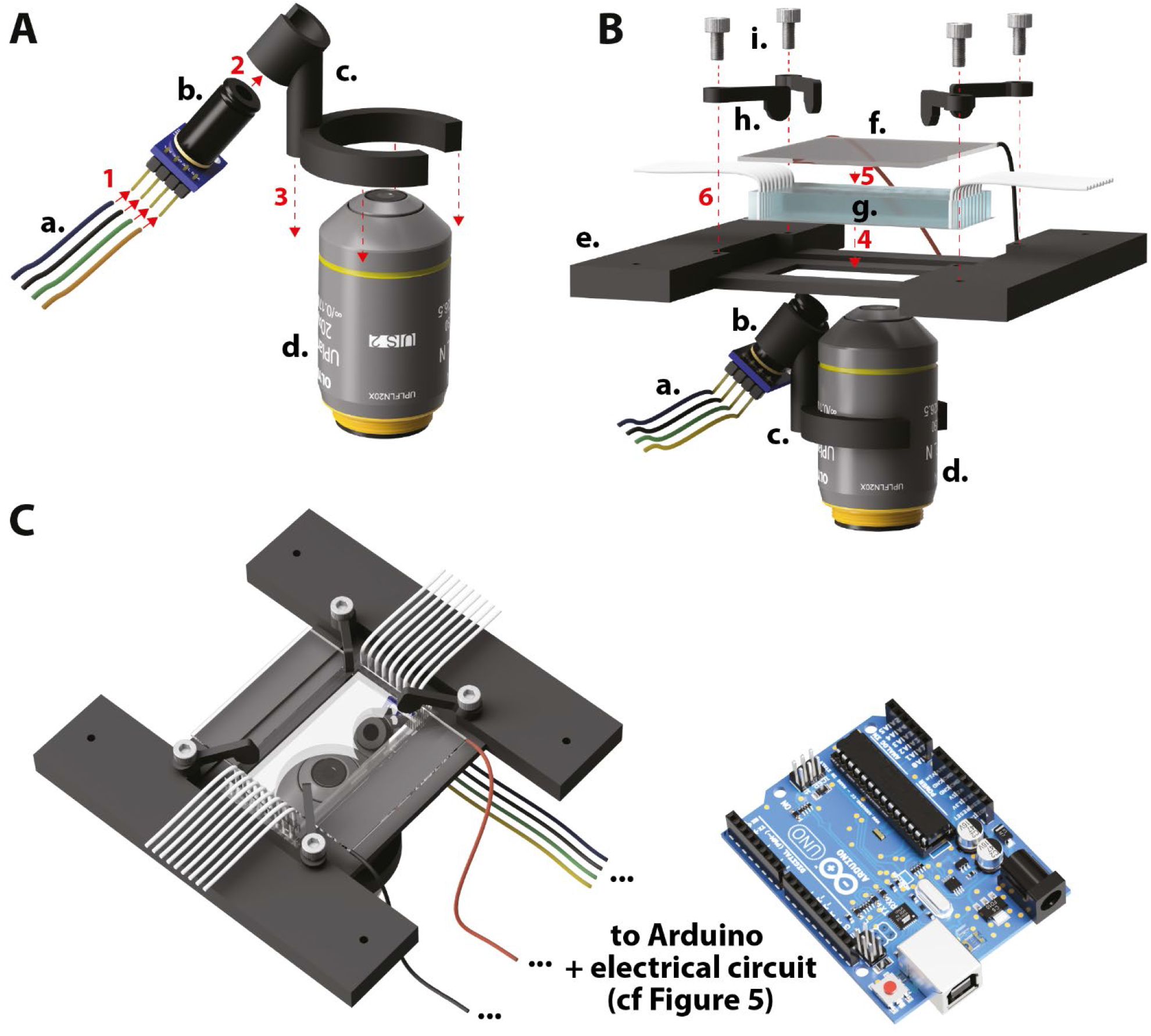
Hardware assembly guide of HeatChips. (A) Step 1: Solder four electrical cables (a) to the four pins of the MLX90614 GY-906 infrared temperature probe (b). Step 2: Insert (b) into the 3D-printed probe holder (c). Step 3: Insert the probe holder around the objective of the inverted microscope (d). (B) Step 4: Place the microfluidic device (e) onto the sample holder (f). Step 5: Place the heating glass (g) onto (e). (Steps 4 and 5 are interchangeable according to the user’s needs and preferences). Step 6: Lock (g) against (e) using 4 M3 screws (h) passing through 4 tightening legs (i) and screwed inside (f). (C) Representative view of the assembled device.

## 6. Operation instructions

To use HeatChips:

1. Install the microfluidic device onto the *sample holder* (Figure 4B, step 1).
2. Put the heating glass in contact with the microfluidic device (Figure 4B, step 2), and gently press on the glass slide of the microfluidic device in order to remove air from the interface and ensure efficient heat transfer.
3. Lock the contact using the four *tightening legs* at the corners of the *sample holder* (Figure 4B, step 3). At this step, the system should look similar to what is depicted in Figure 4C and Figure 5.
4. Switch on the Arduino and plug in the power supply to the micro USB or USB C port.
5. (Optional) If using the *HeatChips_Arduino_Program_withMatlabOutput.ino Arduino program,* run the Matlab plotting program *read_HeatChips.m* to see the temperature of the sample, of the room, and of the power output (PWM) value. If using *HeatChips_Arduino_Program_withMatlabOutput.ino,* you can use any serial reader (such as the built-in monitor in the *Arduino Software*, *cf* code for detailed instructions) to read the Arduino text outputs, indicating the temperature of the sample, of the room, and the power output.
6. Wait until the temperature of the sample reaches equilibrium (5-20min), and run the imaging acquisition.

**Fig. 5.**
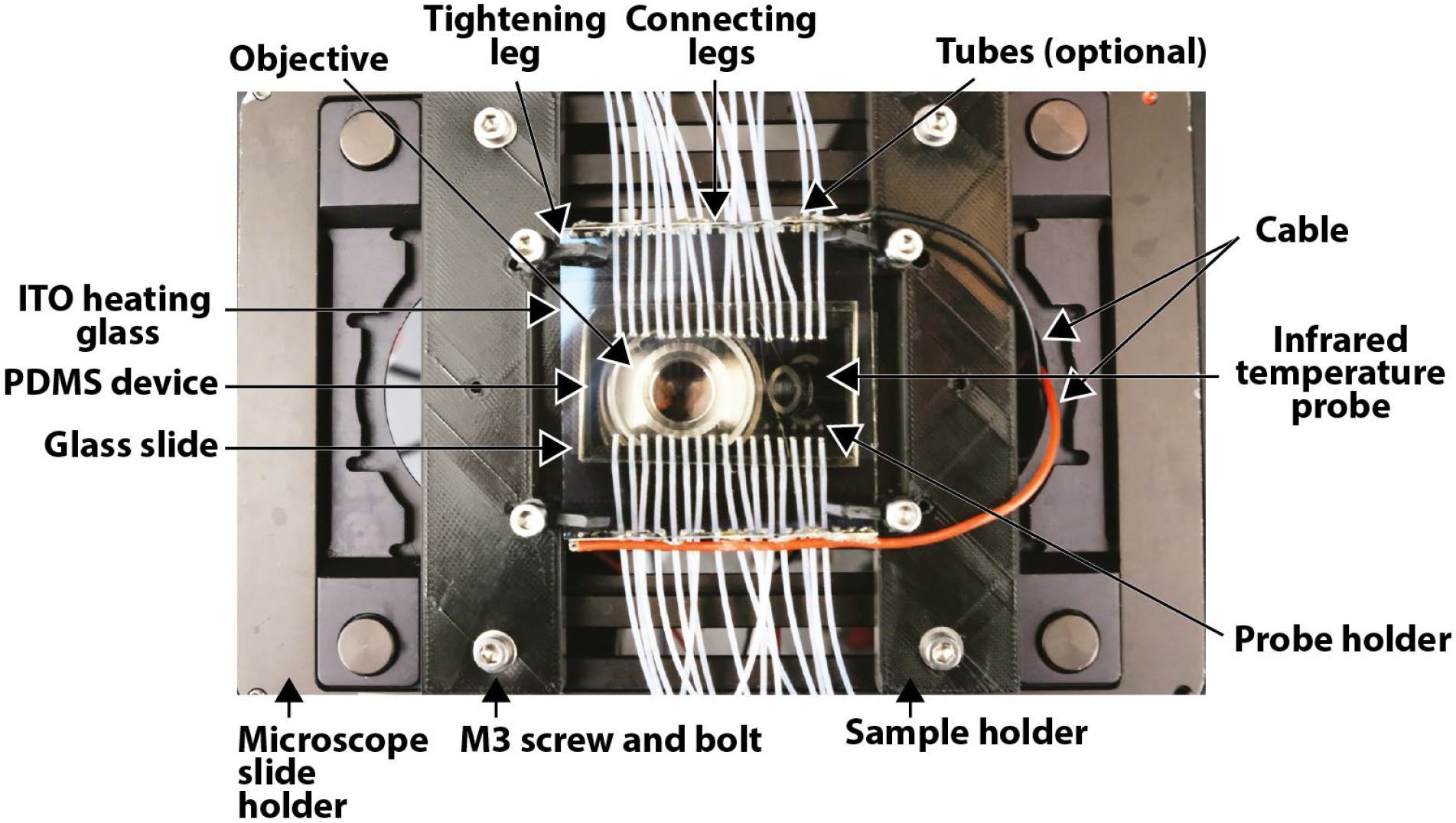
Picture of HeatChips mounted on a microscope stage during a typical experiment, with exhaustive captions.

**Fig. 6.**
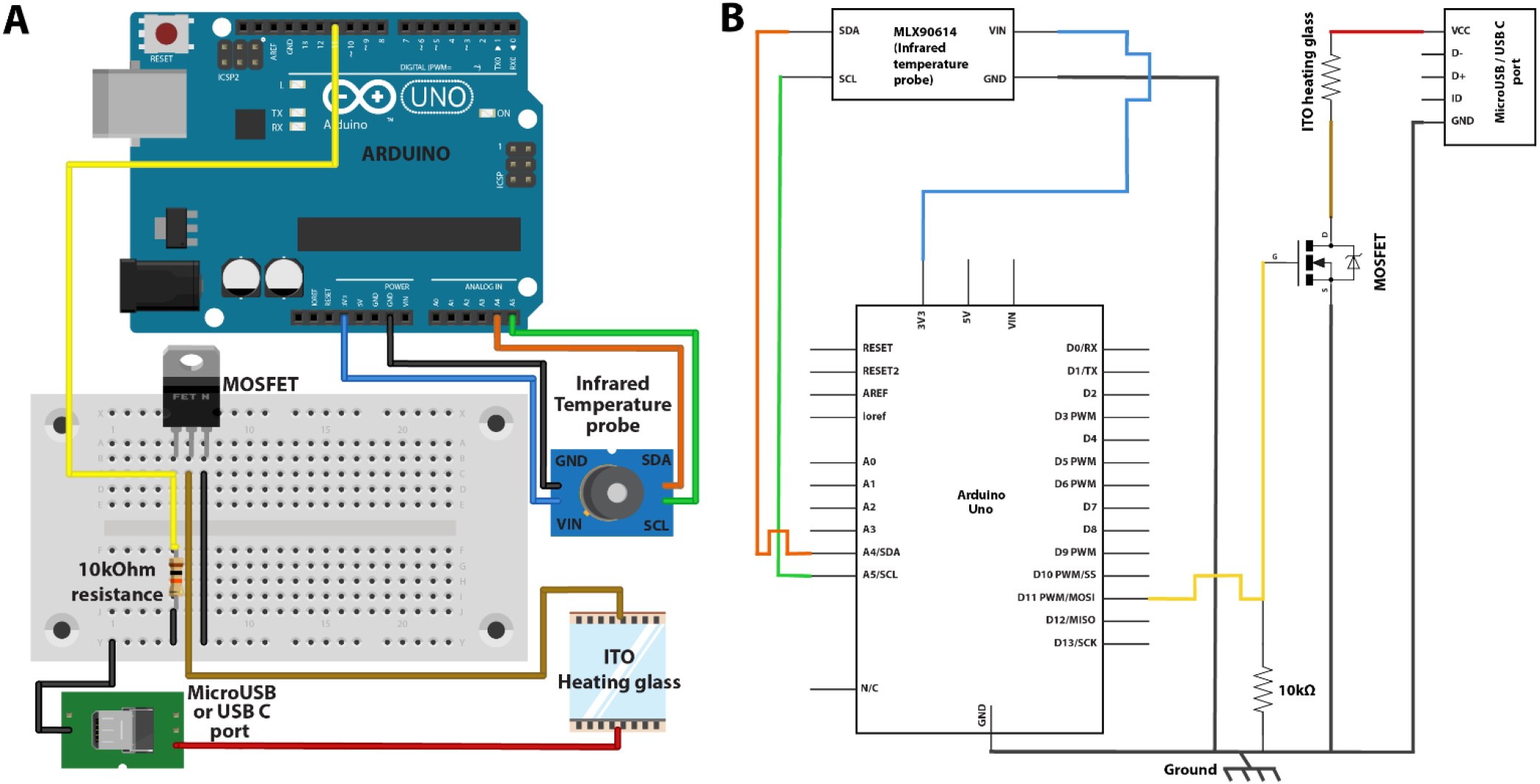
(A) Sketch of the electrical design. The colored lines represent electrical connections. (B) Scheme of the electrical design. The colored lines are the same as in (A).

## 7. Validation and characterization

To validate and characterize the heating system, we inserted a thermistor inside the PDMS device during the casting step. This 2mm thick temperature probe was in contact with the glass slide of the device (Figure 7A). Then, we heated the device using HeatChips, with a setpoint at 30°C, and compared the temperature measured by the infrared probe with that of the thermistor, until equilibrium (Figure 7B, orange lines). In this configuration, the difference in measured temperature was small (2.5%), and can be attributed to the greater depth of measurement of the thermistor. Indeed, there is a temperature gradient between the top of the PDMS block and the glass slide. The probe measures the temperature over its thickness and is inserted inside the PDMS, whereas the infrared probe measures the temperature of the glass slide, which is cooler than the PDMS. Altogether, this confirms that the infrared probe is precisely measuring the temperature of the glass slide, where the sample of interest is located.

In addition, this experiment allowed us to measure the response dynamics of the feedback system. In its configuration (PDMS thickness = 5mm, P coefficient = 50, I coefficient = 10, D coefficient = 0), the 99% equilibrium state was reached after ~12min, with an overshoot of 0.5°C (Figure 7B). The heating speed at full power in this configuration was 1.5°C/min and at equilibrium, and the power used to maintain the sample at 30°C, with a room temperature of 24°C, was 0.4W. These performances are similar to that of heating systems heating the sample from the sample holder with Peltier modules or heating resistors [8]. Of note, modifying the P and I coefficients in the Arduino program (see comments in the code to see the effect of each parameter), or the PDMS thickness, affects the equilibrium time, the heating speed, and the overshoot. For example, it is possible to obtain no overshoot at the cost of a slower heating speed by reducing P, while reducing the PDMS thickness will both increase the heating speed and reduce the overshoot.

To further validate the robustness of the temperature control, HeatChips was used for more than 2h in a room where the temperature was modified by turning on and off the air conditioning (Figure 7C). Under these conditions, a 1.5°C degrees change of the room temperature in 5min led to a change in the temperature of the sample inferior to 0.05°C, which is acceptable for a wide range of microfluidic applications.

HeatChips was also used in routine for published experiments [13,14], to keep yeast cells at 30°C in a 3 to 10mm-thick PDMS microfluidic device, while being imaged every 5min for several days under an inverted microscope, with the room temperature varying between 23 and 26°C. In these experiments, a 10×,20×, or 60× objective was used with brightfield or phase contrast illumination, as well as fluorescence from different wavelengths, without any perturbation from the ITO-coated heating glass.

**Fig. 7.**
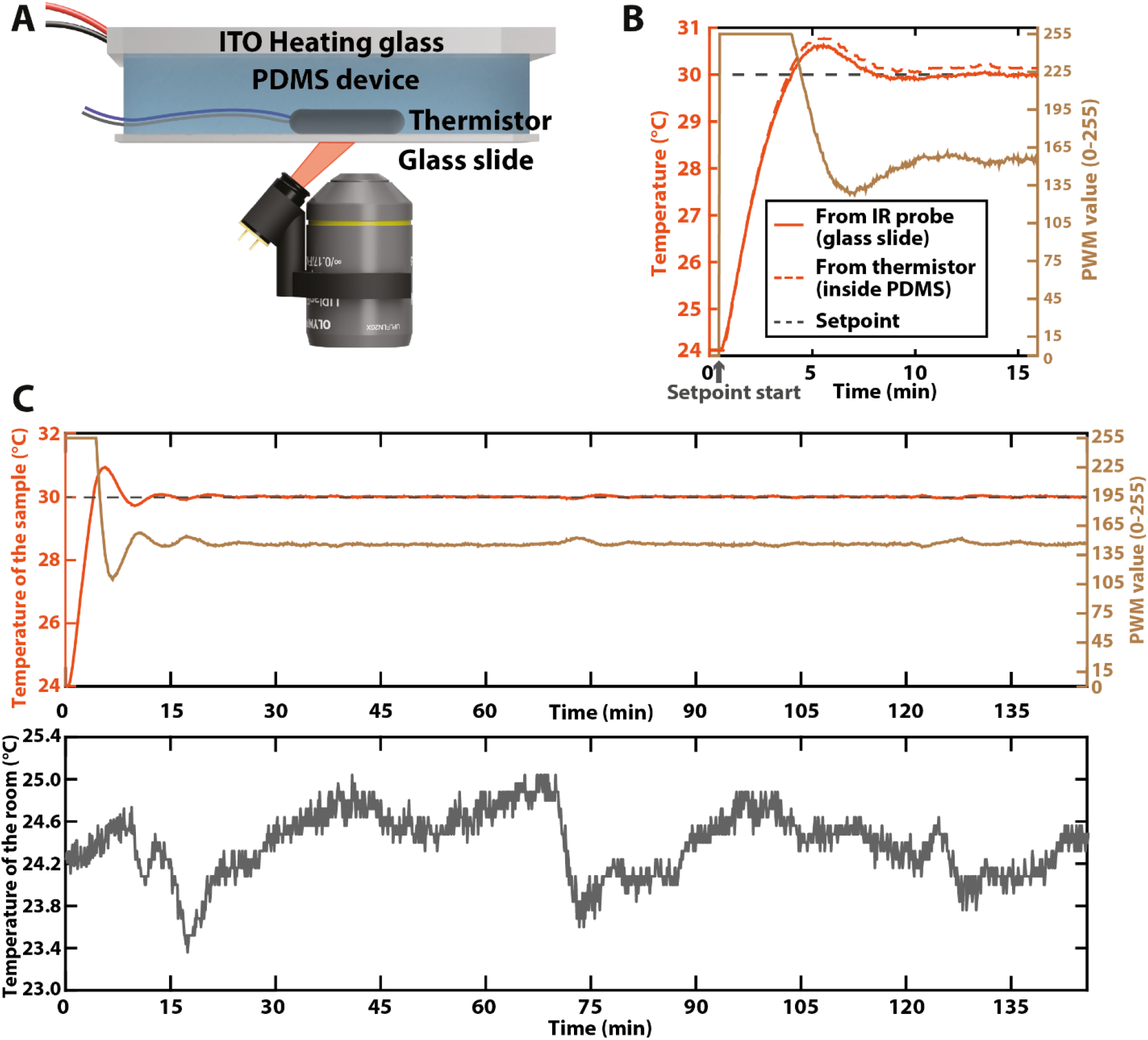
Validation experiments of HeatChips (A) Scheme of the validation experiment using a thermistor temperature probe. The scale of the objects is not representative. (B) Temperature measured by the infrared probe and by the thermistor (orange lines), and PWM output values computed by the control part and sent to the heating part of HeatChips (beige line) after imposing a 30°C setpoint (dashed grey line). (C) (Top) Temperature measured by the infrared probe (orange) and PWM output values computed by the control part and sent to the heating part of HeatChips (beige), with a setpoint value of 30°C (dashed grey line), during an experiment with an unstable room temperature. (Bottom) Temperature of the room associated with the (Top) plot.

## Conclusion

We have presented HeatChips, a microscopy-compatible heating system for microfluidic devices. It is compatible with any PDMS-based microfluidic device and easily adaptable from one microscope to another. In addition, this heating system is small, costs less than 100€, and can easily be reproduced in ~3h (including printing time) by non-expert users following the instructions described in this study. Indeed, it is made only from a heating glass, three 3D printed parts, two electronics components, a phone charger, and an Arduino controlled by a simple Arduino program. HeatChips is especially well-suited for maintaining a constant temperature at the surface of a microfluidic device during long-term time-lapses, and has been extensively used for in-chip monitoring of cell growth for which controlling the gas composition was not required.

## Ethics statements

No ethics statement to report.

## CRediT author statement

**Théo ASPERT:** Conceptualization, Assembling, Testing, Software, Data curation, Writing-Original draft preparation, Writing-Reviewing and Editing.

**Gilles CHARVIN:** Supervision, Writing-Reviewing and Editing, Funding acquisition

## Acknowledgments

We thank Alain Litt for the advice on the electronic design. This work was supported by the PrxAGE - 17-CE11-0034-03 project, a French State fund managed by the Agence Nationale de la Recherche.

## Notes

https://doi.org/10.5281/zenodo.7316523

https://3dprint.nih.gov/discover/3DPX-017678

### Competing Interest Statement

The authors have declared no competing interest.

